## Supplementary Figures for "Motion-corrected eye tracking (MoCET) improves gaze accuracy during visual fMRI experiments"

### Supplementary materials

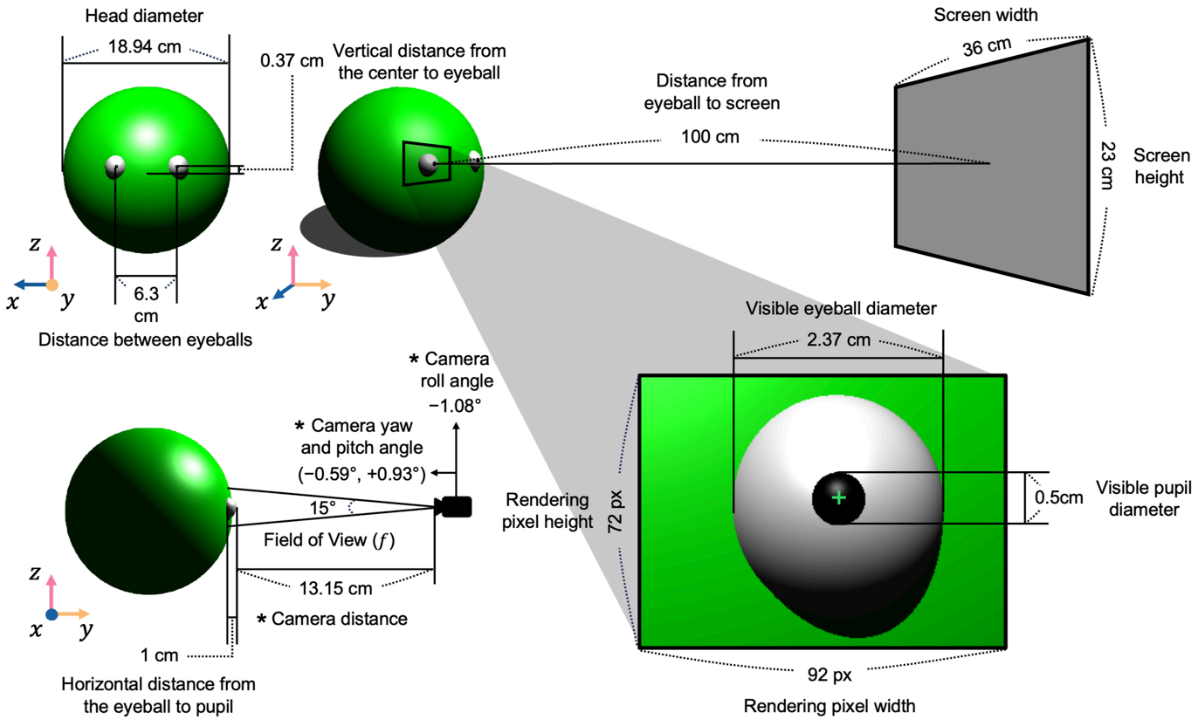


Supplementary Figure 1. Anthropometric parameters used in the model simulation. The head model was designed as an isotropic sphere to simplify head motion dynamics, with the diameter defined by averaging three key human head dimensions: head breadth (ear-to-ear distance), horizontal depth (back of head to nose), and vertical length (chin to top of head), based on measurements from prior studies[29–31]. The model's eyeballs are physically attached to the head sphere, ensuring that head movements directly influence the 3D spatial position of the eyeballs. Each eyeball can independently rotate to simulate gaze shifts, while the pupil, aligned with the gaze direction, is positioned along the line extending from the eyeball center to the target location. To replicate realistic eye tracking conditions, virtual camera parameters—including yaw, pitch, roll, and distance from the eyeball—were customized to match participant-specific eye tracking data (see Supplementary Figure 2 for examples). Averaged values of these camera parameters across participants are indicated with an asterisk (*). For visualization purposes, the example model eye was rendered at a high-resolution of 512×384 pixels, whereas the actual simulation was conducted at 92×72 pixels for computational efficiency.

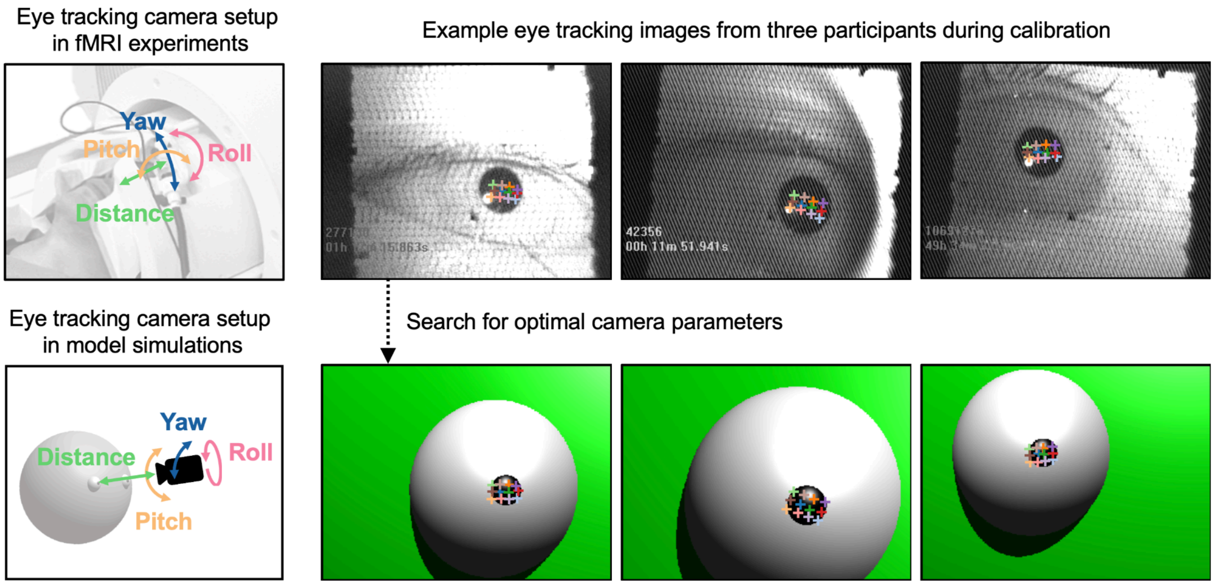


Supplementary Figure 2. Optimization of eye tracking camera parameters in the model simulation. To replicate realistic eye tracking conditions, the virtual eye tracking camera's orientation (yaw, pitch, and roll) and distance from the model's eyeball were systematically adjusted. A total of 36,000 parameters configurations were tested by simulating gaze toward 12 calibration points on the screen. The optimal camera parameters were selected for each individual eye tracking data by minimizing the discrepancy between simulated and actual participant pupil coordinates, ensuring the simulated eye tracking data closely aligned with participant data. The bottom panels show the optimized camera parameters and corresponding simulated pupil locations for 12 calibration points, illustrated for three example participants shown in the top panels.

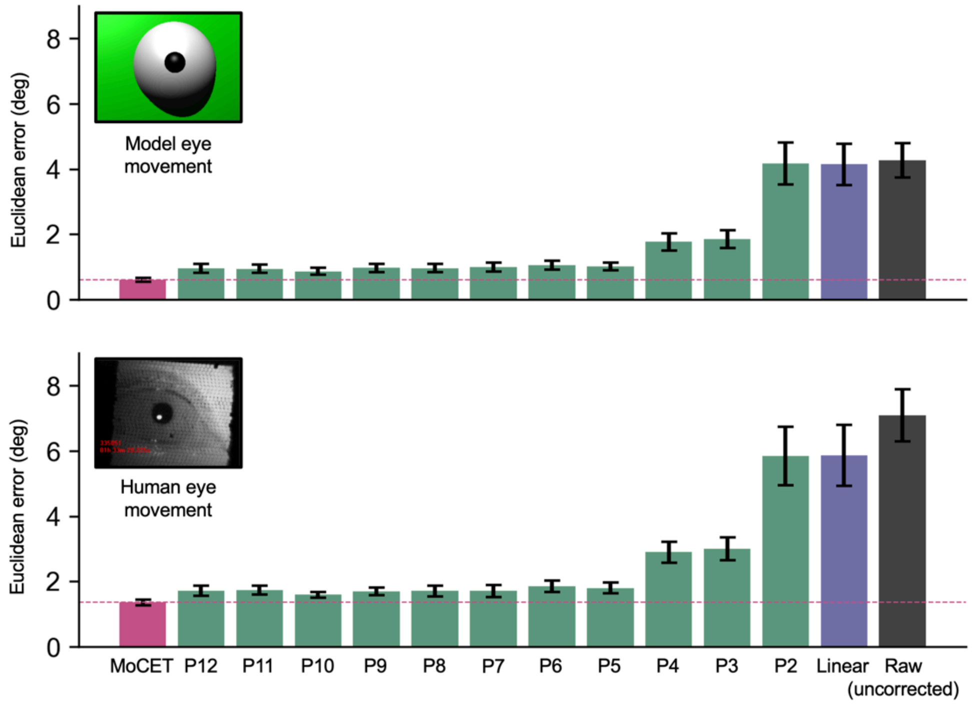


Supplementary Figure 3. Extended analysis of gaze accuracy during the validation stage for model simulation (top) and human eye tracking data (bottom) from Figure 4A. Gaze accuracy is compared across MoCET and various detrending methods, including linear detrending and polynomial detrending up to the 12th order. Polynomial methods show diminishing returns in performance improvement beyond the 5th order, while MoCET consistently achieves the highest accuracy across both datasets. Horizontal dashed lines indicate MoCET's performance as a benchmark. Error bars represent the standard error across participants.

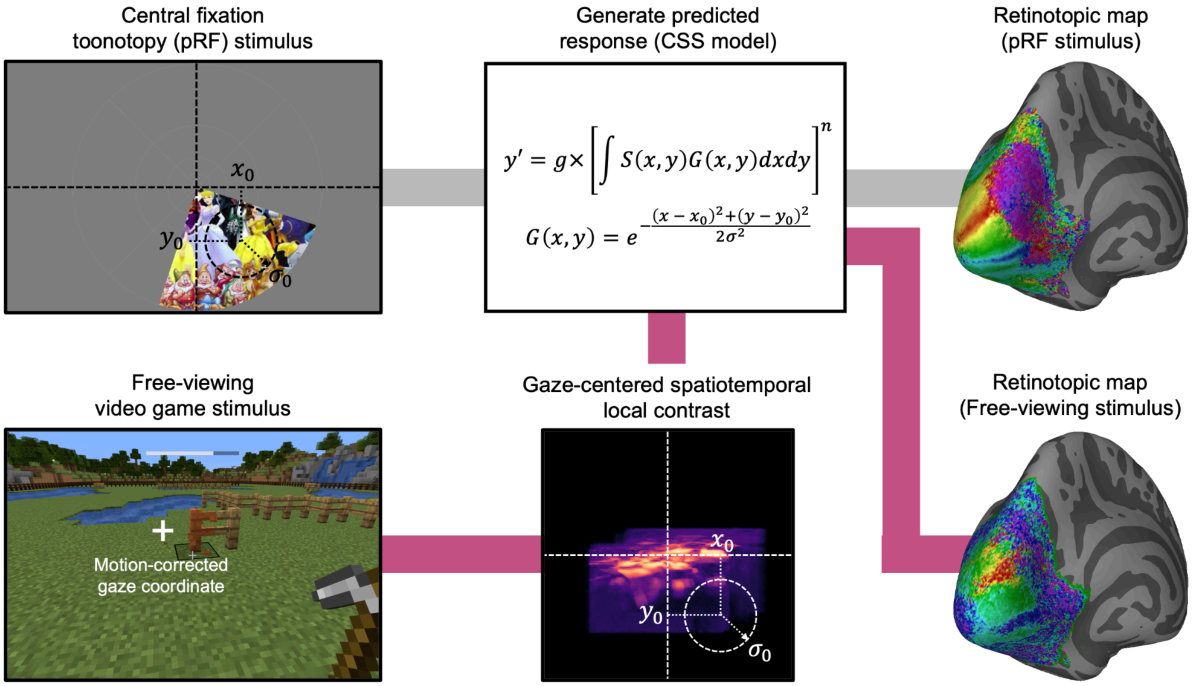


Supplementary Figure 4. Estimation of retinotopic mapping from pRF experiments and free-viewing video game stimuli. Retinotopic mappings in early visual areas were derived using structured pRF stimuli with central fixation (top) and free-viewing video game stimuli (bottom). For pRF experiments, structured stimuli[51] such as wedges, rings, and bars were presented while participants maintained central fixation. Neural responses were modeled using the Compressive Spatial Summation (CSS) model[44] to estimate retinotopic map across the visual cortex. For free-viewing experiments, detrended eye tracking data (e.g., MoCET or polynomial detrending) was used to transform visual stimuli into gaze-centered spatiotemporal local contrast maps[42], which served as input to the pRF model.

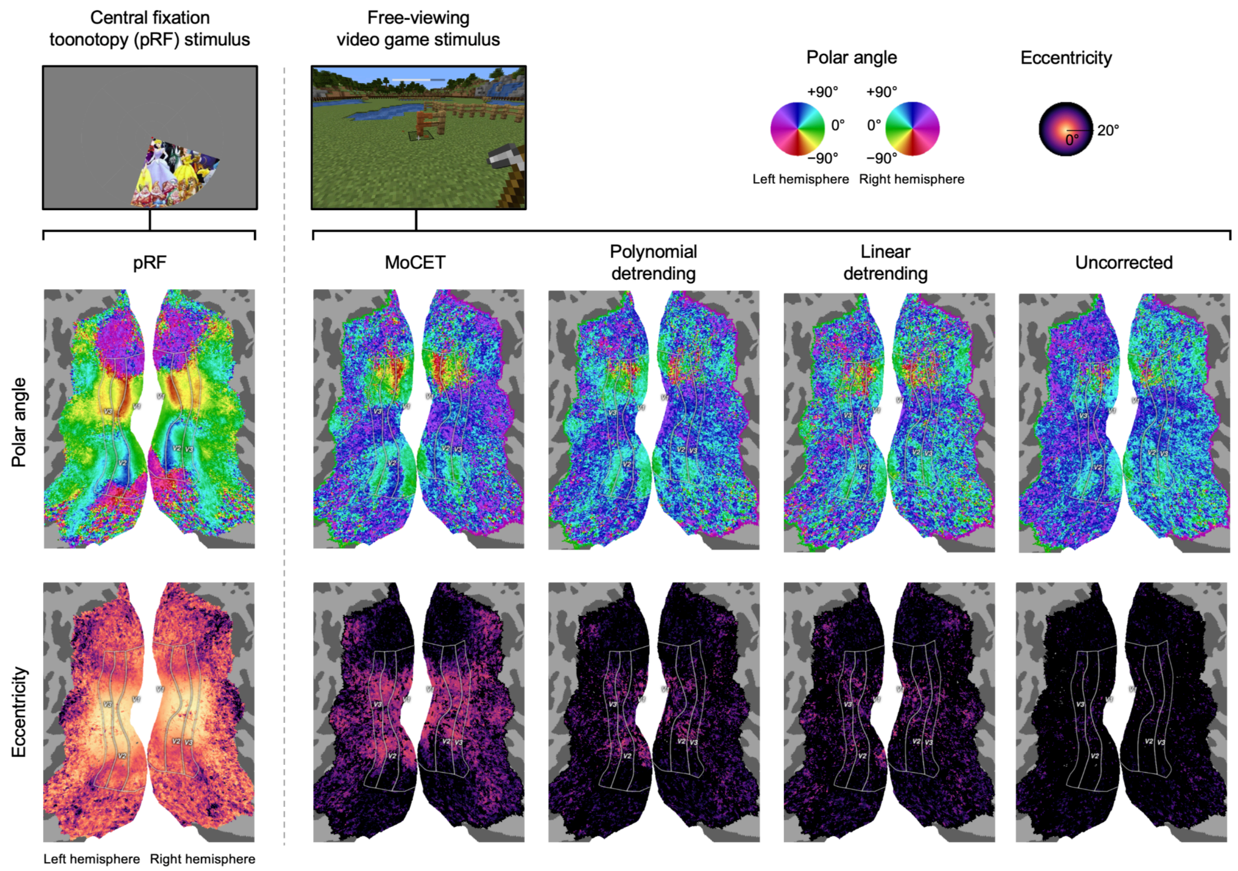


Supplementary Figure 5. Group-averaged retinotopic maps estimated from pRF experiments and free-viewing visual stimuli. Flattened cortical maps display polar angle retinotopy in early visual areas (V1, V2, V3) for both hemispheres. Visual field maps are derived using structured pRF experiments with central fixation (pRF) and free-viewing video game stimuli corrected with MoCET, polynomial detrending, linear detrending, or uncorrected data. Regions of interest (ROIs) for V1, V2, and V3 were defined based on the Human Connectome Project (HCP) retinotopy dataset[52].
